# Holomycin from *Yersinia ruckeri* mediates fish infection at low temperature

**DOI:** 10.64898/2026.04.29.721558

**Authors:** Yaxin Wu, Jiaquan Huang, Tingjie Zhan, Jun Wu, Jiaxuan Li, Qiandi Gao, Haiming Xu, Xiaojie Li, Yibin Yang, Song Meng, Junya Wang, Yi Yu, Jun Zou, Hai Deng, Heqian Zhang, Zhiwei Qin

## Abstract

Why microbes invest energy in producing structurally complex natural products at extremely low concentrations within hosts remains an important topic of interest in microbial ecology. This is because such compounds with potential “biomolecular activities” can reflect molecule-target interactions, thereby providing conceptual and mechanistic insights that inform the rational design of essential clinical agents. Here, we uncover the *in situ* ecological function of the dithiolopyrrolone antibiotic holomycin produced by the fish pathogen *Yersinia ruckeri*. We show that holomycin’s biosynthesis is strictly gated by low temperature and that trace amounts of holomycin accumulate *in vivo* during infection in rainbow trout. In addition, the holomycin-deficient mutant leads to markedly reduced virulence. Integrated multiomics, phenotypic analyses, and *in vivo* bioassay test reveal that holomycin is not a direct lethal factor but instead remodels host-associated microbiota and enhances *Y. ruckeri* biofilm formation. These coordinated effects promote pathogen colonization and infection progression. Our findings establish holomycin as a host-associated effector and illustrate how its existence and ecological function are mechanistically intertwined in the evolution of pathogenic bacterial chemical arsenals.

## Introduction

Microbial pathogens inhabit densely populated host-associated ecosystems in which successful infection depends on direct interactions with the host, as well as on competition, cooperation, and chemical communication with surrounding microbial communities. Increasing evidence suggests that specialized secondary metabolites (or natural products) from microbes are central to these processes, functioning as conditional effectors that reshape local ecological landscapes during infection^1-3^. Yet a fundamental question remains unresolved: how do such metabolites exert profound biological effects *in vivo* despite often (if not always) being produced only transiently and at extremely low concentrations^4^? Addressing this question requires a combined understanding of their intrinsic bioactivities, together with the environmental signals and regulatory programs that govern when and where they are deployed.

*Yersinia ruckeri* is one of the most economically important bacterial pathogens in cold-water aquaculture and the causative agent of Enteric Redmouth Disease (ERM), which affects salmonids and other cultured fish worldwide^5-7^. In addition, despite its rarity, a clinical report has described the isolation of *Y. ruckeri* from a human wound infection following aquatic exposure, raising concerns regarding the zoonotic potential^8^. In the case of fish infection, a striking and long-recognized feature of this pathogen is its strong temperature dependence: disease outbreaks are favored under cooler conditions (approximately 16-18 °C), whereas virulence is markedly reduced or even abolished at higher temperatures^9^. Although classical determinants such as secretion systems, iron acquisition pathways, and extracellular toxins contribute to pathogenesis, they do not adequately explain this sharply gated low-temperature infection phenotype^10-12^. Therefore, *Y. ruckeri* may rely on environmentally responsive strategies beyond canonical virulence factors to establish infection in cold-water hosts.

We previously found that *Y. ruckeri* produces the small-molecule natural product holomycin, a representative member of the dithiopyrrolones (DTP) natural product family^13^. DTPs possess a unique bicyclic pyrrolinodithiole scaffold and exhibit broad-spectrum antibacterial and antitumor activities (Figure 1A)^14-16^. The mode of action of DTPs was initially attributed mainly to selective inhibition of RNA synthesis or interference with carbohydrate metabolism during cellular respiration^17-20^. More recent studies, however, have shown that DTPs can chelate zinc ions and thereby inhibit multiple JAMM-domain-containing deubiquitinases, providing new insights into their molecular targets and mechanisms of bioactivity^21-23^.

**Figure 1.**
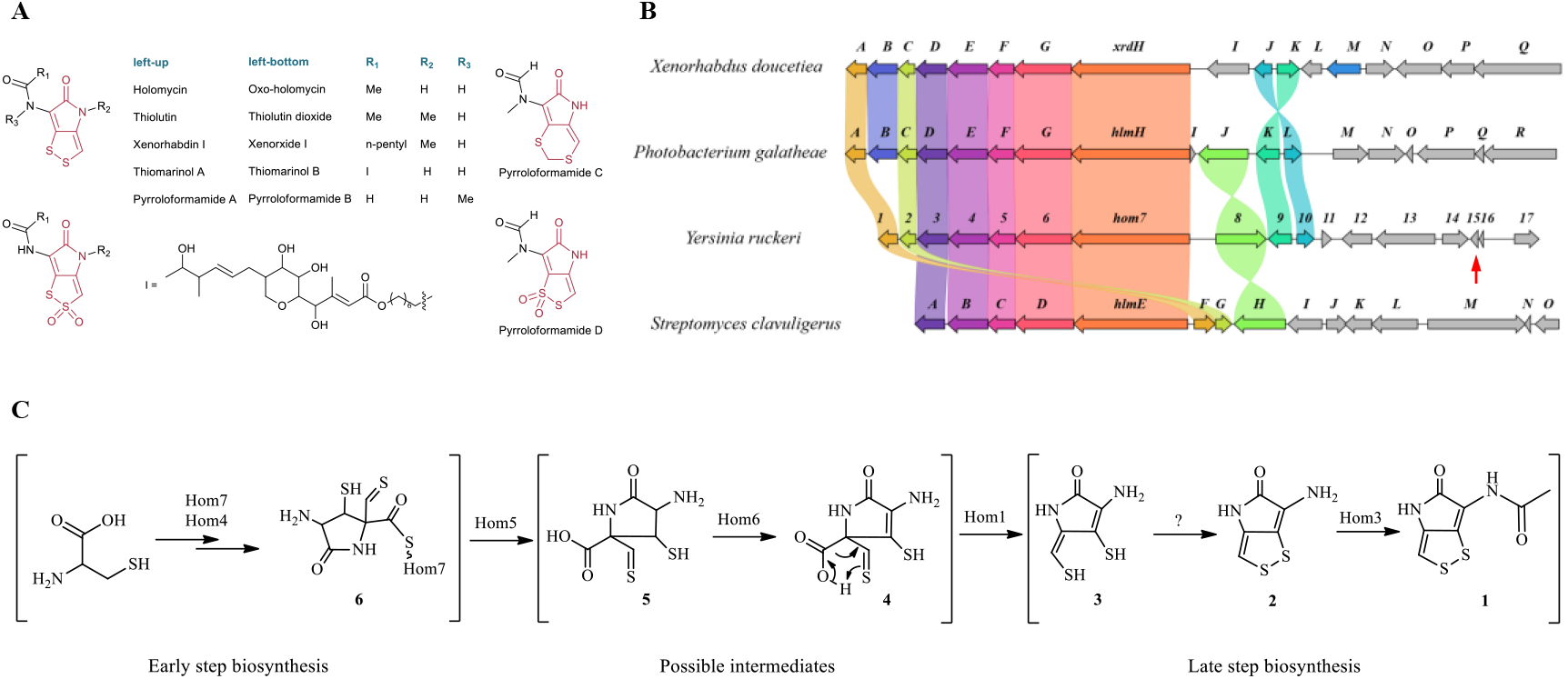
Dithiolopyrrolone natural products and holomycin biosynthesis. (A) Representative chemical structures of dithiolopyrrolone natural products, including holomycin, thiolutin, aureothricin, and thiomarinol. (B) Comparison of holomycin BGCs highlights the presence of a unique cold shock-like protein, Hom15, in *Y. ruckeri*. (C) The proposed biosynthetic pathway of holomycin in *Y. ruckeri*.

The *in situ* ecological functions of DTPs in natural environments have not been systematically investigated, and current understanding is limited to only a few isolated examples. For instance, holomycin has been shown to promote biofilm formation in the marine bacterium *Photobacterium galatheae*, providing direct experimental evidence for its potential ecological role in environmental adaptation and host colonization^24^. However, in the case of *Y. ruckeri* infection, it is unknown whether holomycin is synthesized during infection, whether its production is specifically triggered by low temperature, and whether it contributes directly to pathogenic progression. Resolving these questions is important not only for understanding the unusual temperature-dependent pathogenicity of *Y. ruckeri*, but also for clarifying how bacterial pathogens deploy specialized metabolites within host-associated environments.

In our previous and recent unpublished work, through the comparative analyses of holomycin biosynthetic gene clusters (BGCs) from diverse microbial sources, we identified a unique cold-shock protein-encoding gene, *hom15*, present exclusively in the *Y. ruckeri* cluster (Figure 1B). Subsequent multi-omics analyses revealed that transcription of *hom15* is highly coordinated with key holomycin biosynthetic genes. Moreover, fermentation experiments confirmed that holomycin production is strictly dependent on low-temperature conditions. Our previous work further demonstrated that Hom15 plays a crucial regulatory role in maintaining normal holomycin biosynthesis at low temperature, and deletion of *hom15* significantly reduces product accumulation under such conditions^13^. These findings suggest that Hom15 may ensure efficient operation of the holomycin pathway by maintaining the stability and translational efficiency of biosynthesis-related transcripts in cold environments, thereby functionally coupling temperature sensing with secondary metabolic regulation. In addition, it is speculated that holomycin is unlikely to be a fortuitous antibacterial by-product; rather, it may represent a functional small molecule specifically activated under precise temperature control, operating through mechanisms distinct from those of classical toxins.

Here, we combine bioinformatics, chemical profiling, and *in vivo* cold-temperature infection models to define the ecological role of holomycin during *Y. ruckeri* infection in fish. We show that holomycin biosynthesis is strictly activated at low temperature, accumulates *in vivo* at trace levels during infection, and is indispensable for full pathogenicity in rainbow trout. Unexpectedly, holomycin is not a direct lethal toxin to the host. Instead, it functions as an ecological interference molecule: it reshapes the host-associated microbiota and promotes *Y. ruckeri* biofilm formation at varying concentrations, thereby facilitating colonization, immune evasion, and infection progression. Our work reveals holomycin as a host-associated effector whose biosynthetic specialization and ecological function are mechanistically intertwined, providing a framework for understanding how pathogenic bacteria deploy specialized metabolites to manipulate microbial communities and host environments during disease.

## Results

### Phylogenetic analysis of the holomycin biosynthesis-related gene cluster from different producers

To investigate whether the holomycin produced by *Y. ruckeri* contributes to this thermal-dependent pathogenicity, we first analyzed the whole-genome sequence of a highly virulent *Y. ruckeri* strain isolated from Channel catfish (*Ictalurus punctatus*) infected with ERM. Genome mining revealed a 12-kb holomycin biosynthesis-related gene cluster (*hom* BGC), similar to our previous work^13^. While holomycin biosynthesis has been extensively characterized in gram-positive *Streptomyces clavuligerus*^21,25^, our previous comparative analysis revealed that the *Y. ruckeri hom* BGC exhibits a markedly different organizational logic (Figure 1B and 1C). A striking evolutionary distinction is the presence of a CSP encoding gene, *hom15*, physically integrated within the *Y. ruckeri hom* cluster. Deletion of *hom15* significantly reduces the production of holomycin^13^. Phylogenetic analysis demonstrated that the *hom* BGC forms a distinct clade that remains phylogenetically isolated from traditional actinomycete producers. Within this clade, the *hom*-associated CSP is conserved across pathogenic *Y. ruckeri* and entirely absent in all other known holomycin BGCs (Figure 1B). The tight physical coupling of a cold-responsive regulator within the *hom* BGC provides a strong evolutionary rationale for the thermal gating of holomycin production in *Y. ruckeri*, suggesting an adaptive production strategy during the low-temperature pathogenic program.

To directly assess the temperature dependence of holomycin biosynthesis, we cultured *Y. ruckeri* across a series of temperatures (16, 18, 20, and 28°C). As shown in Figure 2A, holomycin was robustly produced at 16 and 18°C, with the highest levels observed at 16 °C; in contrast, no detectable production occurred at 20 or 28 °C. These results indicate a negative correlation between culture temperature and holomycin output. To determine whether this thermal constraint is unique to *Y. ruckeri*, we conducted an extensive survey of the literature describing holomycin production in other microbial hosts ^24,26-34^. Notably, all reported fermentation conditions for non-*Y. ruckeri* producers were conducted above 20 °C, typically at 28 or 30 °C, consistent with their growth optima.

**Figure 2.**
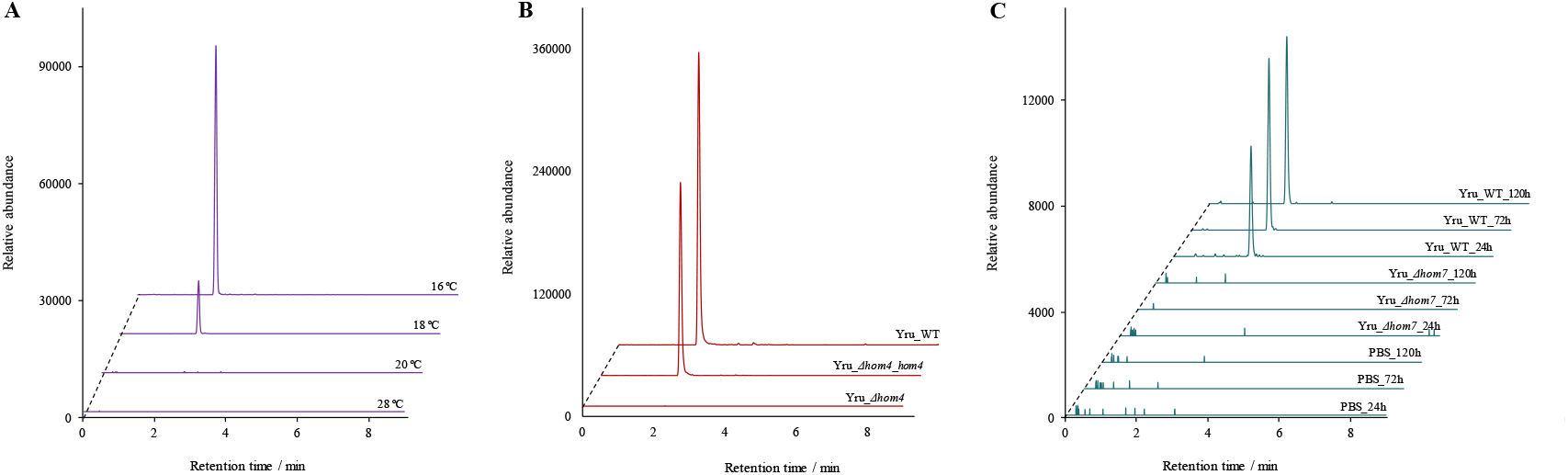
Holomycin production in *Y. ruckeri*. (A) Mutational analysis of genes required for holomycin biosynthesis. Production was abolished in Yru_*Δhom7* and Yru_*Δhom4* and restored in the complemented strains. (B) In vivo detection of holomycin during infection. Trace amounts of holomycin accumulated in rainbow trout injected with Yru_WT at 24, 72 and 120 h post-infection, whereas no holomycin was detected in Yru_*Δhom7-* or PBS-treated fish. (C) Temperature-dependent production of holomycin in *Y. ruckeri*. Holomycin was produced at 16 °C and 18 °C but not at 20 °C or 28 °C.

Taken together, these findings demonstrate that holomycin biosynthesis in *Y. ruckeri* is specifically activated under low-temperature conditions. This temperature-dependent regulation suggests a potential ecological role for holomycin during cold-associated infection within the host.

### Multiomics profiling of the essential genes for holomycin biosynthesis in *Y. ruckeri*

To fully characterize the *hom* BGC in *Y. ruckeri*, we employed an integrative multiomics approach to gain deeper insights into gene function and regulation in this gram-negative strain. To this end, *Y. ruckeri* was cultured in two different media at 16 °C: tryptic soy broth (TSB) medium, which supports robust holomycin production (6.4 mg/L), and de Man-Rogosa-Sharpe (MRS) medium, which does not support detectable production. We hypothesized that genes involved in holomycin biosynthesis would exhibit significant expression changes under holomycin-producing conditions. To test this hypothesis, we performed combined transcriptomic and proteomic analyses using cell biomass collected 48 h after cultivation in each medium (hereafter referred to as the TSB group and MRS group). This time point was selected on the basis of prior time-course high-performance liquid chromatography (HPLC) analysis showing that holomycin biosynthesis in *Y. ruckeri* typically peaks within 24 h, ensuring that sufficient transcriptomic and proteomic signatures are detectable by 48 h. Preliminary differential expression analysis revealed 1641 differentially expressed genes (DEGs) whose |log_2_FC| > 1 and 547 differentially expressed proteins (DEPs) whose |log_2_FC| > 1 in the TSB group. Notably, several genes associated with holomycin biosynthesis, including *hom1-7* and *hom15*, exhibited such expression patterns (Figure 3A and 3B). The overall downregulation of *hom* genes at this stage does not contradict pathway activation, as secondary metabolite biosynthesis is often regulated through temporally coordinated transcriptional programs^35^. To further investigate the regulatory dynamics, we examined the expression levels of *hom1-7* by qRT-PCR at multiple time points (6-96 h). The results revealed that these *hom* genes were significantly upregulated at early stages and reached a plateau at approximately 48 h under cultivation at 16 °C, followed by gradual downregulation at later time points (Figure 3C). Therefore, the apparent transcriptional repression at this later stage is consistent with feedback regulation and pathway shutdown following enzyme accumulation and metabolite buildup, a phenomenon widely observed for toxic or energetically expensive secondary metabolites^36^. Collectively, these findings demonstrate that the *hom* BGC and its integrated CSP operate as a synchronized functional unit under low temperature. This pathway follows a temporal pattern in which holomycin biosynthesis undergoes an early activation phase followed by a late repression phase rather than continuous induction throughout the fermentation period.

**Figure 3.**
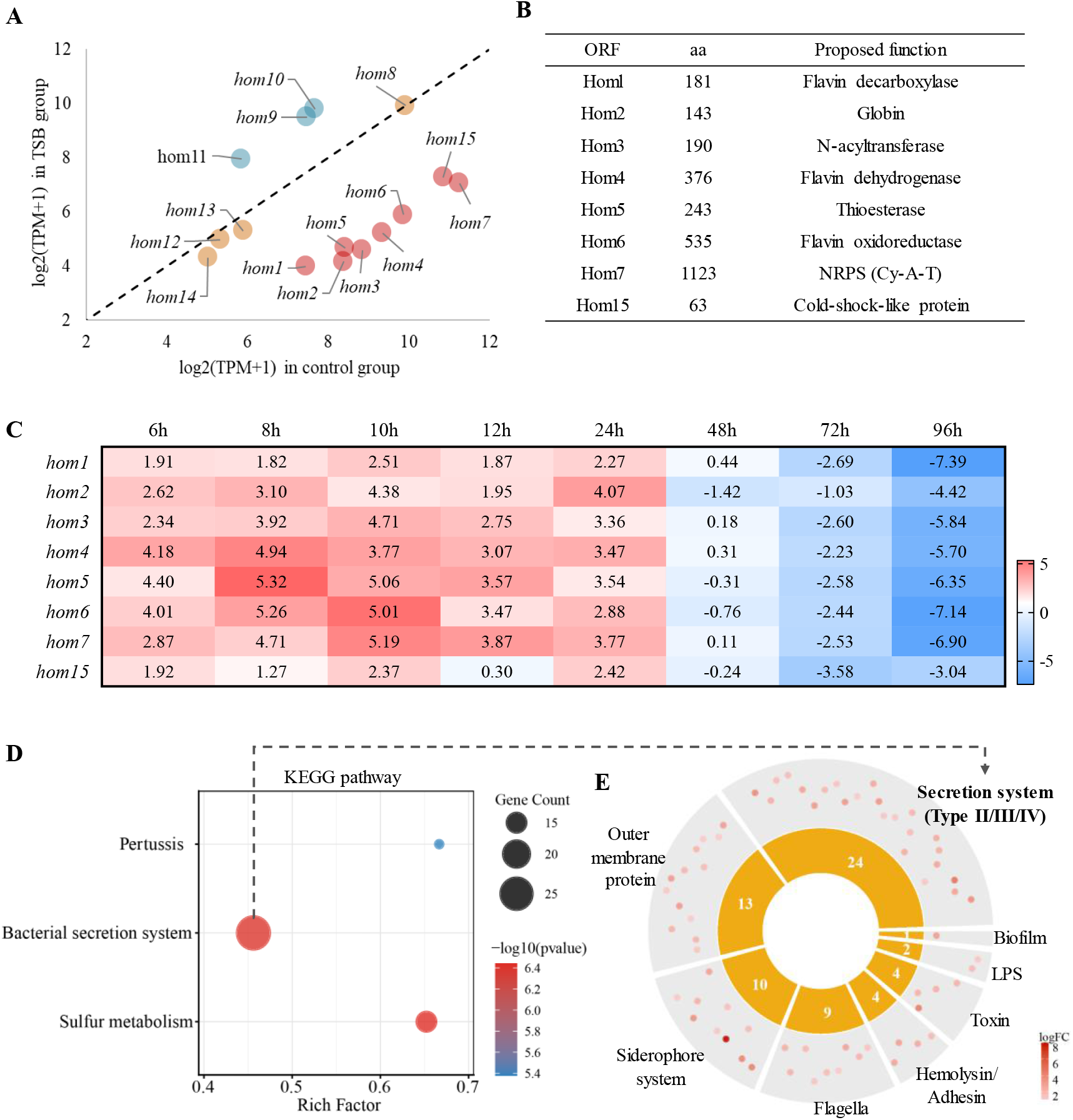
Transcriptomic profiling of *Y. ruckeri* in TSB (holomycin-producing) relative to the control. (A) Comparison of the expressions of holomycin biosynthetic genes (*hom1*-*hom7* and *hom15*) between TSB and MRS groups. Data are shown as log_2_(TPM+1) values. The dashed line indicates equal expression. Genes below the dashed line are expressed at lower levels in the TSB. (B) Number of aa and proposed functions of Hom1-Hom7 and Hom15. (C) Heatmap of the relative mRNA expression of *hom1*-*7* and *hom15* in *Y. ruckeri* from 6 h to 96 h. The colours indicate the relative expression levels. (D) KEGG pathway terms for the upregulated DEGs. (E) Catergory of virulence factors with fold change regulated by *Y. ruckeri* cultured in TSB medium compared to MRS medium.

KEGG enrichment analysis of the DEGs revealed that upregulated transcription profiles were mainly enriched in the bacterial secretion system and pertussis (Figure 3D), which are closely associated with the expression of virulence factors. The transcriptome sequence of *Y. ruckeri* was annotated using Prokka and RAST. As shown in Figure 3E, nine families of virulence factor-encoding genes were identified, including Type II/III/IV secretion system, outer membrane proteins, siderophore system, flagella, hemolysin, adhesin, LPS, biofilm, and other toxins. *Y. ruckeri* cultured in TSB significantly upregulated these virulence factors encoding gene compared to the MRS group, suggesting that holomycin biosynthesis in *Y. ruckeri* plays an important role in regulating the expression of its virulence factors.

Finally, for multiple purposes in this study, as described in the following sections, we constructed in-frame deletion mutants in *hom7*, generating the strain Yru_*Δhom7*. We selected *hom7* as the target because, as suggested in Figure 1C, Hom7 is predicted to function at an early stage of holomycin biosynthesis. Its deletion therefore eliminates the formation of any pathway-specific intermediate that might otherwise confound our infection assays (see following sections). Consistent with our previous findings, the production of holomycin was completely abolished in the mutant strain Yru_*Δhom7* but could be restored by ectopic expression of the corresponding gene (Figure 2B). Notably, the deletion of *hom7* did not cause any significant alteration on the ultrastruction of *Y. ruckeri* (Figure S1).

### *In vivo* chemical profiling confirms the thermal-gated accumulation of holomycin during fish infection

To determine whether holomycin is a synchronized component of the *Y. ruckeri* infection process rather than a laboratory artefact, we employed a rainbow trout (*O. mykiss*) infection model at 16 °C. Fish were intraperitoneally injected with the wild - type strain (Yru_WT) at a bacterial dose of 50 µL (OD_600_ = 0.1) or with phosphate-buffered saline (PBS) as a negative control.

Using our established liquid chromatography high-resolution mass spectrometry (LCMS) method, we sought to determine whether holomycin is genuinely produced during this pathogenic process. Detection of whole-fish extracts at 72 h post-infection revealed a clear holomycin signal exclusively in samples from Yru_WT-infected rainbow trout, which displayed retention times and MS features identical to those of the authentic standard (Figure 2B, Figure S2-S3). Quantification analysis further revealed that holomycin accumulates rapidly in the early stages, reaching 0.20 µg/fish at 24 h and plateauing at approximately 0.33 µg/fish by 72-120 h (Figure 2B). To the best of our knowledge, this is the first study to provide direct *in vivo* chemical evidence that holomycin is actively produced and accumulates during infection by the wild-type pathogen. LCMS further revealed that no holomycin was detected in fish infected with the Yru_ *Δhom7* mutant (Figure 2C).

Together, above results suggested that *Y. ruckeri* has repurposed a core cold-stress component to serve as a molecular trigger, potentially linking environmental sensing to the deployment of its specialized metabolites during the pathogenic program. If this hypothesis is correct, then it will raise fundamental considerations regarding the biological role of holomycin in the fish host during *Y. ruckeri* infection. This question will be investigated in greater detail in the following sections.

### Holomycin is indispensable for *Y. ruckeri* infection but is not a direct lethal factor

To investigate the pathogenic role of holomycin during the interaction between *Y. ruckeri* and its fish host, rainbow trout fish were intraperitoneally injected with Yru_WT, Yru_*Δhom7*, Yru_*Δhom7* supplemented with holomycin (3 µg/fish), and holomycin alone (3 µg/fish) all at a bacterial dose of 50 µL (OD_600_ = 0.1) (Figure 4A). A PBS-treated group served as a negative control. Then a 14-day time course survival assay showed distinct infection dynamics between the different groups (Figure 4B). Yru_WT-infected fish exhibited a statistically significant reduction in survival and reached 0% survival rate by day 12. In contrast, the virulence of fish treated with Yru_*Δhom7* markedly attenuated, with survival rate of 73% by day 12. Compared with the mutation alone, supplementation of Yru_*Δhom7* with holomycin partially restored pathogenicity, leading to a significant decrease in survival rate (38%); however, the resulting survival rate remained significantly higher than that caused by Yru_WT. Notably, compared with the PBS control, the administration of holomycin alone did not lead to a significant difference in survival, indicating that holomycin alone does not cause mortality in rainbow trout.

**Figure 4.**
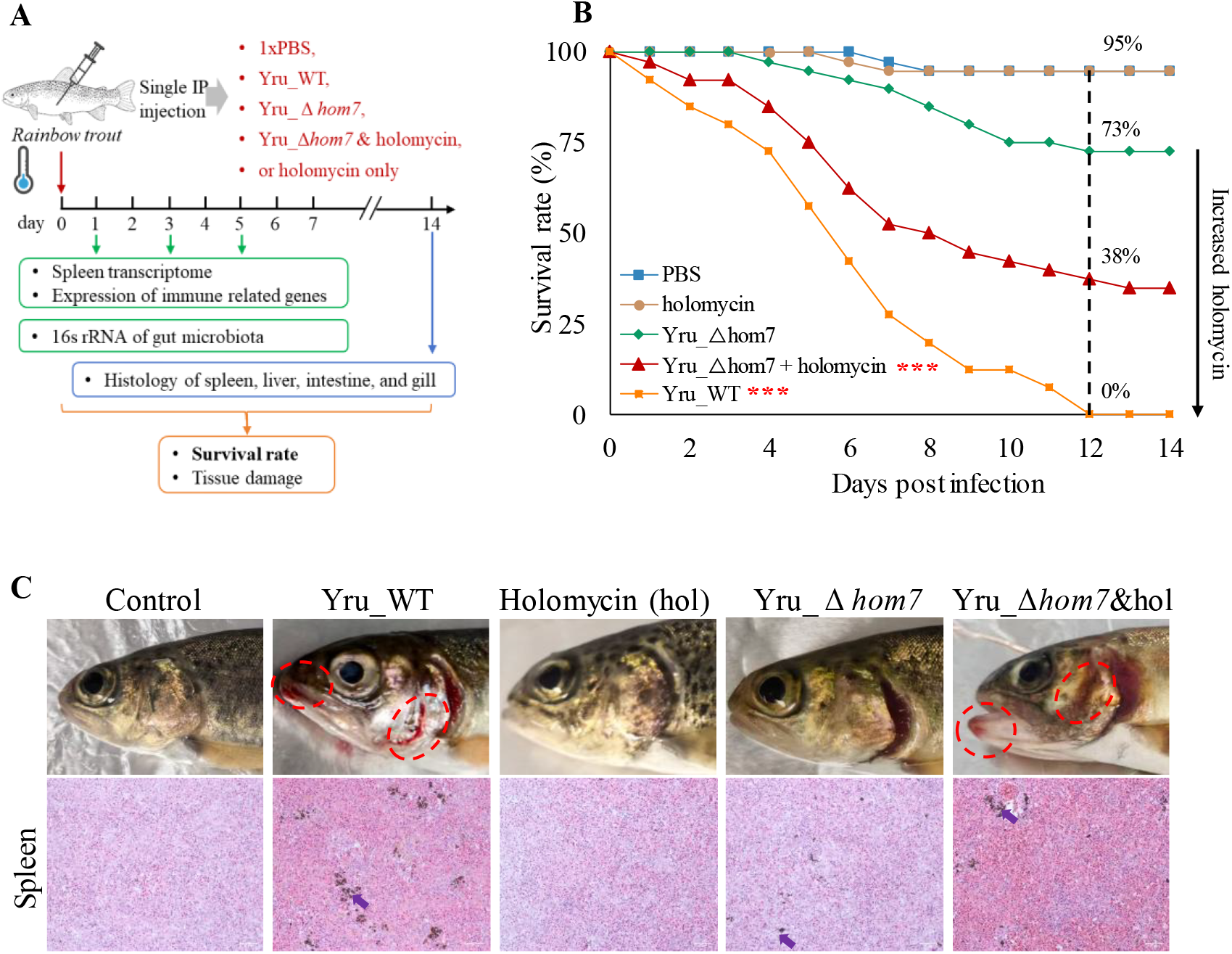
Holomycin promoted *Y. ruckeri* infection. (A) Schematic of *Y. ruckeri* infection in rainbow trout. (B) Survival rate curves of rainbow trout over 14 days following a single intraperitoneal injection of PBS, holomycin alone (3 μg), Yru_WT, Yru_Δ*hom7*, or Yru_Δ*hom7* supplemented with holomycin. (C) Representative H&E-stained spleen sections from from fish in each group at 72 h post-infection. Purple arrows indicate melanomacrophage centers. * represents the comparison of treatment groups to control group (PBS). Survival curves were analyzed using a two-way analysis of variance with Tukey’s multiple comparisons test. \**P* < 0.05, \*\**P* < 0.01, and \*\*\**P* < 0.001.

Histological examination was also performed on spleen, a major immune organ in fish to assess infection-associated pathology. As shown in Figure 4C, red pulp expansion with erythrocyte engorgement was observed in Yru_WT and Yru_*Δhom7*_holomycin infected groups, accompanied by the formation of multiple melanomacrophage centers (MMCs), the histological markers of immune and infection response. Fish infected with Yru_*Δhom7* only exhibited mild splenic changes compared with control, while there was no significantly alteration in the holomycin-treated spleen. Similarly, the liver, intestine, and gill also displayed histological damage in Yru_WT and Yru_*Δhom7*_holomycin, but not in the PBS, holomycin alone, or Yru_*Δhom7* groups (Figure S4). These data demonstrate that holomycin is required for the *Y. ruckeri*-induced tissue pathology, consistent with its role as a key virulence determinant in this infection model (Figure 4C). Above infection-associated observations were further supported by the increased transcription levels of inflammatory factors, including *il-1β, il-6, il-8* and *il-10*. As shown in Figure 5A, the mRNA expression of the above biomarkers was dramatically elevated in spleen of Yru_WT-infected fish, increasing by 30-fold, respectively, in comparison to those in the control group. In Yru_*Δhom7*-infected fish, which did not accumulate holomycin *in vivo*, only a slight induction of those inflammatory genes was induced.

**Figure 5.**
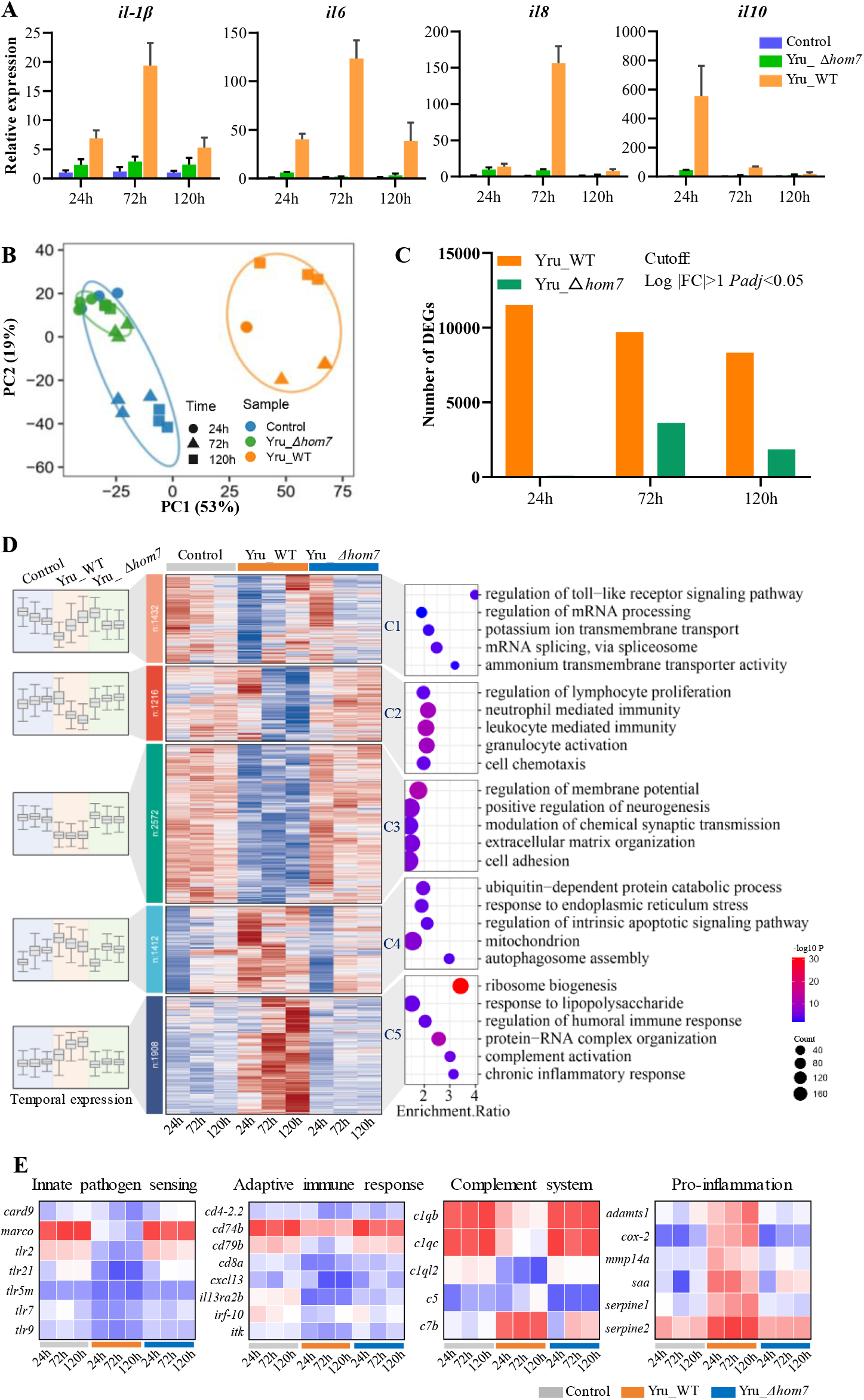
Transcriptome profiling of the spleen of rainbow trout showing potential holomycin interruption. (A) Relative mRNA expression of *il-1β* in the spleen at 24 h, 72 h, and 120 h post-infection. Data are shown as the mean ± SD (N = 6). (B) Principal component analysis (PCA) of the transcriptome in the spleens of fish infected with PBS, Yru_WT, and Yru_Δ*hom7* at 24 h, 72 h, and 120 h post-infection. Shape and colour indicate the time point and group, respectively. (C) Numbers of DEGs altered by Yru_WT and Yru_Δ*hom7* compared with the control. (D) Time-course clustering of splenic DEGs (C1-C5) at 24 h, 72 h, and 120 h, shown as z scored expression heatmaps with corresponding temporal expression patterns (left, boxplots) and GO enrichment results (right, bubble charts). (E) Expression levels of innate and adaptive immune genes in PBS-, Yru_WT-, and Yru_Δ*hom7*-infected fish at 24 h, 72 h, and 120 h post-infection. Data are shown as the mean ± SD and were analysed with Student’s t test; \**P* < 0.05, \*\**P* < 0.01, and \*\*\**P* < 0.001.

### Holomycin is required for *Y. ruckeri* to suppress host immune responses

To better understand the mechanistic role of holomycin during *Y. ruckeri* infection, we performed a whole transcriptomic analysis of the spleen, collected from trout infected with Yru_WT or Yru_*Δhom7*, alongside uninfected controls (treated with PBS) at 24, 72, and 120 h post-infection. Both principal component analysis (PCA) and sample-to-sample correlation analysis based on VST-normalized expression matrices revealed that Yru_WT infection group were clearly separated from the other two groups, forming an independent cluster. The first two principal components together explained approximately 69% of the total variance (PC1: 49%, PC2: 20%) (Figure 5B). In contrast, samples from the control group and the Yru_*Δhom7* infection group clustered together in the PC1-PC2 space. At each time point, the two groups largely overlapped or were positioned in close proximity, with no obvious separation. At the transcriptome level, Yru_WT triggered immediate and potent transcriptional alteration in fish spleens, reaching 11524 DEGs at 24 h and remaining high at 72 h (9704 DEGs) and 120 h (8336 DEGs). Yru_*Δhom7* showed a delayed and weaker response, with minimal perturbation at 24 h, with only 92 DEGs, followed by a later increase at 72 h (3636 DEGs) and a slight decrease to 1855 DEGs at 120 h (Figure 5C). Temporal trajectory analyses further indicated a holomycin-dependent shift in the timing and potency of the splenic response to infection (Figure 5D). DEGs were grouped into five clusters across three time points, each displaying a unique temporal profile. Notably, genes in Yru_*Δhom7* and control groups exhibited similar expression patterns, while Yru_WT infection induced markedly distinct temporal trajectories.

Downregulated clusters C1 and C2 were characterized by a rapid and delayed suppression of DEGs, respectively, in Yru_WT infected fish. Both clusters were enriched for innate immune processes including neutrophil activation and TLR signalling, indicating that *Y. ruckeri* rapidly inhibited early innate defences in a holomycin-dependent manner. C2 were also enriched for regulation of lymphocyte proliferation, suggesting that suppression extends to the initiation of adaptive immune priming. Cluster C3 exhibited consistently downregulated expression across three time points by Yru_WT, with enrichment in membrane potential and extracellular matrix organization. Among upregulated clusters, C4 showed early induction of DEGs in Yru_WT infected fish, and were enriched for pathways associated with ubiquitin-dependent protein catabolic process and acute cellular stress responses in endoplasmic reticulum/mitochondrial. C5 showed progressive upregulation over the infection time. and was enriched for sustained inflammation-related pathways. Consistent with these clustering results, transcription levels of immune related genes (Figure 5E), including innate pathogen sensing, adaptive immune response, were decreased by Yru_WT compared to Yru_*Δhom7*. Complement C1q recognition complex (*c1qb, c1qc, c1ql2*) were also downregulated, while downstream components (*c5* and *c7b*) and several pro-inflammatory mediators, such as *cox-2* and *saa*, were significantly upregulated. Importantly, the absence of holomycin attenuated the transcriptional changes across all five clusters, directly implicating its role in splenic immune reprogramming driven by *Y. ruckeri*.

Overall, the findings in this section indicate that holomycin is a key virulence determinant contributing to *Y. ruckeri* pathogenicity. Its presence is associated with a complex host immune state during *Y. ruckeri* infection, characterized by the suppression of host immune recognition and adaptive immune-associated signals alongside sustained chronic inflammatory activation.

### Holomycin promotes *Y. ruckeri* colonization by reshaping host microbiota and enhancing biofilm formation

For decades, holomycin has primarily been studied as a broad-spectrum antibiotic. Its defensive role in *Y. ruckeri* infection suggests that holomycin may function as a microbiome-targeting effector that destabilizes or reorganizes the microbial community within infected tissues. To test this hypothesis, we further assessed the pathogenicity of the Yru_WT and Yru_*Δhom7* strains in rainbow trout by analysing the composition and abundance of microbiota within the fish. To this end, 16S rRNA sequencing was performed. At the community level, α-diversity (Chao1 richness and Shannon diversity) and *β*-diversity (unweighted UniFrac) were evaluated among the groups at 24, 72, and 120 h (Figure 6A). In terms of *α*-diversity, Chao1 richness increased in both infection groups compared with that in the PBS group at 72-120 h, whereas Shannon diversity significantly increased earlier in the Yru_WT group than in the Yru_*Δhom7* group. *β*-Diversity (Figure 6B and Table S1) showed significantly consistent separation of the Yru_WT group from the PBS group at 24-120 h (*R*^*2*^ = 0.192-0.243), whereas the Yru_*Δhom7* group exhibited weaker and decreased separation (significant only at 72 h; *R*^*2*^ = 0.186), suggesting that holomycin deficiency attenuated *Y. ruckeri* -associated microbiota restructuring. Taxonomic profiling further revealed pronounced compositional shifts at the phylum level in the Yru_WT group relative to those in the PBS group, characterized by a significant increase in the abundance ratio of *Bacteroidota* and *Firmicutes*, accompanied by a marked increase in the abundance of *Proteobacteria* (Figure S5), while in the Yru_*Δhom7*-treated fish, the abundance of those phylum was slightly affected. At the genus level, the abundance of *Yersinia*, was undetectable in the PBS control but was consistently detected in Yru_WT-infected fish (Figure 6C), whose absolute abundance increased from 217.8 at 24 h to 698 at 72 h and 484 at 120 h, whereas its abundance remained lower in the Yru_*Δhom7* group (Figure 6D). Compared to the PBS control, Yru_WT-treated fish displayed decreased abundances of *Muribaculum* (24-120h), a gut probiotic for immune regulation ^37,38^, and increased abundances of *Cetobacterium* at 24h, a probiotic for gut defenses^39^, while Yru_*Δhom7*-induced weaker effects on these gut probiotics (Figure 6D). Taken together, *Y. ruckeri*, when producing holomycin, can significantly reshape the gut microbiota structure, composition and abundance during infection in fish.

**Figure 6.**
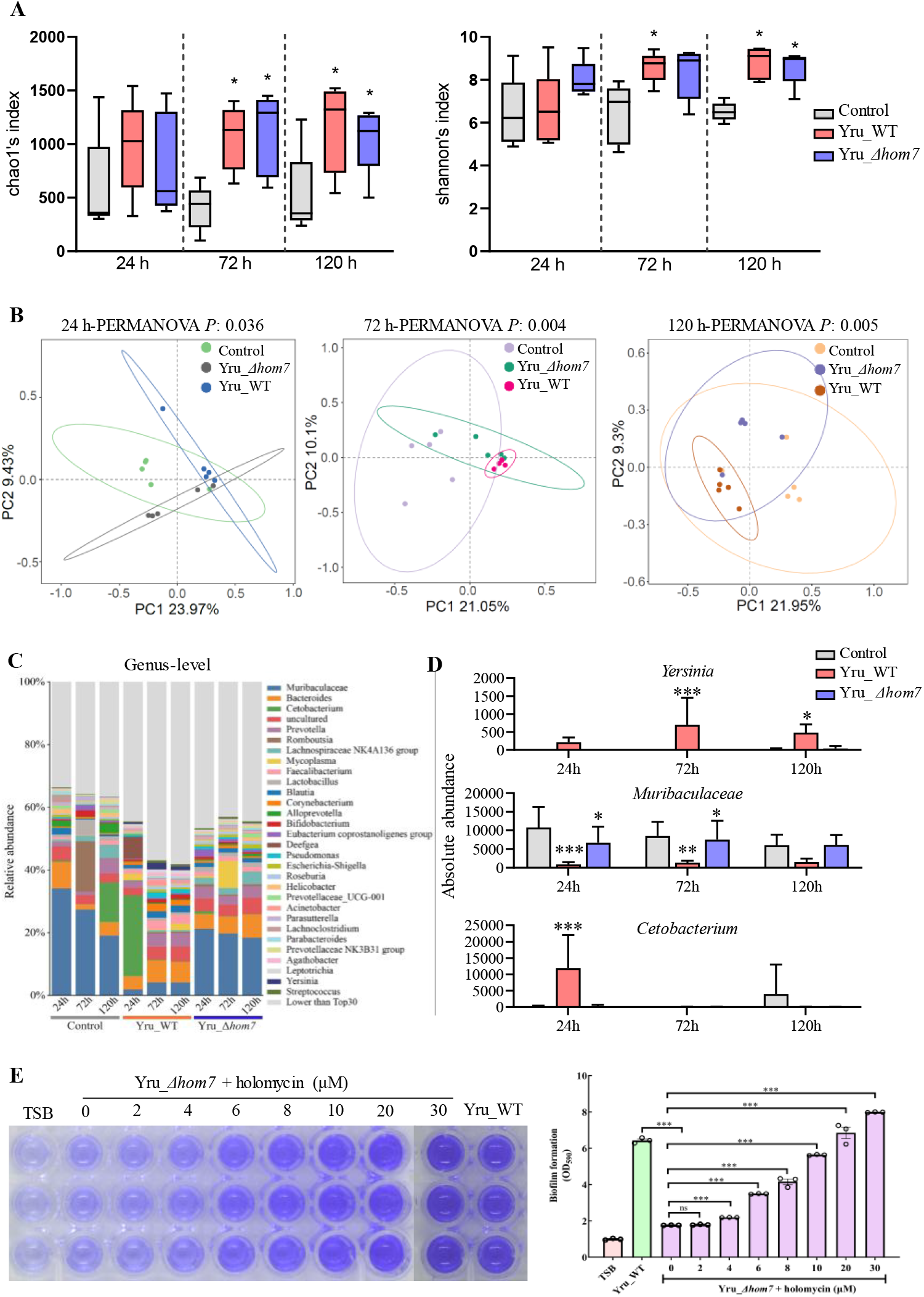
Holomycin-associated microbiome remodelling and dose-dependent promotion of biofilm formation. (A) α-diversity of the gut microbiota at 24, 72, and 120 h post-infection, shown as Chao1 richness (left) and Shannon diversity (right). Data are shown as the mean ± SD and were analysed with the Kruskal-Wallis test and pairwise Wilcoxon test. (B) *β*-diversity of the gut microbiota at 24, 72, and 120 h post-infection visualized by PCoA based on unweighted UniFrac distance. Data were analysed with PERMANOVA. (C) Genus-level composition of rainbow trout gut microbiota at 72 h post-infection. Top 30 genus are displayed. The remaining taxa are grouped as “Lower than Top30”. (D) Absolute abundance of *Yersinia, Muribaculaceae* and *Cetobacterium* in fish infected with PBS, Yru_WT, or Yru_Δ*hom7* at 24 h, 72 h, and 120 h post-infection. (E) Representative crystal violet staining of biofilms formed by Yru_*Δhom7* supplemented with holomycin (0-30 µM) (left panel) after 72h of incubation. Control and Yru_WT were the negative and positive controls, respectively. Quantification of biofilm biomass relative to that of the TSB control (right panel). Data are shown as the mean ± SD and were analysed by Two-way ANOVA; \**P* < 0.05, \*\**P* < 0.01, and \*\*\**P* < 0.001.

Successful infection by *Y. ruckeri* usually depends on effective colonization of fish tissues, a process reminiscent of bacterial biofilm formation. A recent study has reported that holomycin can increase biofilm formation in its native producer, the gram-negative marine bacterium *Photobacterium galatheae*^24^. Biofilms are central to microbial ecology and pathogenesis and can substantially promote bacterial colonization on host tissues or environmental surfaces^40^. For example, in several major pathogens such as *Vibrio cholerae*, biofilms function as cooperative infection strongholds: beyond serving as a protective extracellular matrix, they can prime invading cells through elevated expression of virulence determinants, including toxin-coregulated pilus (TCP) and cholera toxin (CTX), thereby enabling bacteria to enter the host in a pre-armed state^41^. Biofilms also provide an efficient platform for antagonistic systems such as the type VI secretion system (T6SS), while matrix components simultaneously shield resident cells from counterattack^42^. Moreover, their three-dimensional architecture can reshape spatial colonization patterns on host surfaces, optimizing tissue attachment and invasion efficiency^43^. On this basis, we next sought to investigate whether holomycin exerts a similar biofilm-enhancing effect in *Y. ruckeri*. To this end, the biofilm formation was subsequently quantified in the wild type strain Yru_WT, the holomycin-deficient mutant Yru_*Δhom7*, and the holomycin-rescued Yru_*Δhom7* supplemented with various concentrations of purified holomycin. Crystal violet staining assays revealed that Yru_WT formed robust biofilms, whereas the Yru_*Δhom7* mutant exhibited markedly reduced surface-associated biomass. Strikingly, exogenous addition of holomycin restored biofilm formation to wild-type levels when applied at concentrations between 10 and 20 μM (Figure 6E). These results demonstrate that holomycin promotes biofilm development in *Y. ruckeri*, thereby enhancing the pathogen colonization in host during infection.

## Discussion

The ecological functions of microbial secondary metabolites depend critically on their mode, timing, and quantitative levels of production within natural environments. However, substantial differences between laboratory cultivation and actual infection settings often cause the titres, regulatory patterns, and even biological activities observed *in vitro* to diverge markedly from those that occur *in vivo*, thereby obscuring the true ecological roles of many natural products. In this work, we integrate *in vivo* detection, host transcriptomics, infection model with gut microbiome profiling to define the role of holomycin in *Y. ruckeri* infection. Our results demonstrate that (i) holomycin is actively produced during *Y. ruckeri* infection and is required for full virulence in this model, yet it alone did not induce infection responses in fish; (ii) holomycin play an essential role in facilitating pathogen colonization and virulence through coordinated modulation of the microbial community and biofilm formation. A possible model has been proposed in Figure 7.

**Figure 7.**
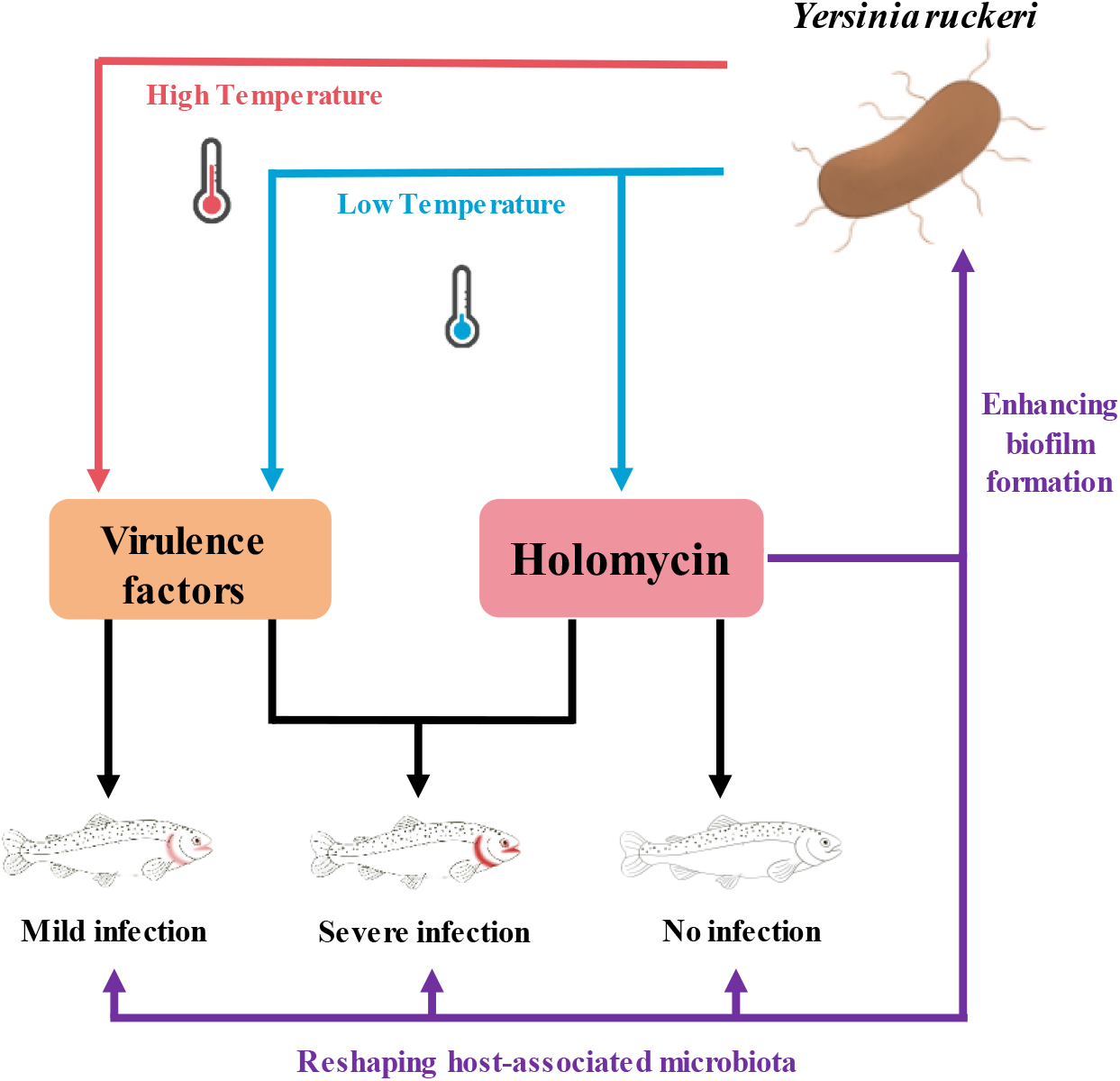
A proposed model for Yersinia ruckeri cold-temperature infection in fish mediated by holomycin. Holomycin functions as a host-associated ecological effector by reshaping host microbiota and enhancing biofilm formation, thereby promoting colonization, immune evasion, and infection progression in cold-water fish. This model illustrates how a specialized metabolite can serve as a temperature-gated virulence strategy linking environmental sensing, microbial competition, and pathogenic success.

Microbial secondary metabolites can mediate chemical competition within host-associated microbial communities in response to stress cues. Such chemical warfare has been documented in leafcutter ant, where the pathogen *Escovopsis* upregulates specialized metabolites to suppress protective microbial partners and weaken host defenses^44^. Holomycin has been linked to stress-responsive regulation^24,45^ and exerts antibacterial effects by chelating metal ions^21^, yet its chemical role in host-pathogen interaction has remained unclear. Here, we show that holomycin accumulates in *Y. ruckeri*-infected fish under cold-temperature conditions (Figure 2), accompanied by stronger microbiome restructuring in Yru_WT than in the holomycin-deficient Yru_*Δhom7* (Figure 6). At the phylum level, Yru_WT infection was associated with enrichment of *Proteobacteria* and depletion of *Bacteroidota* and *Firmicutes*, a community configuration commonly associated with infection-related dysbiosis^46-48^ (Figure S5). This pattern was accompanied by taxon-level differences, including a marked reduction in *Muribaculaceae* and a transient increase in *Cetobacterium* at 24 h in the Yru_WT group (Figure 6C and 6D). Because *Muribaculaceae* have been linked to carbohydrate and nutrient metabolism in other vertebrates, their depletion may indicate loss of commensal functions relevant to community stability, although this remains to be validated in fish^49^. Likewise, the transient increase in *Cetobacterium*, a common fish gut genus associated with enhanced resistance to pathogen infection, may represent an early ecological response to infection, although its functional significance in this context remains unclear^39^. Together with the broad-spectrum antimicrobial holomycin, these observations support the idea that holomycin functions as an “ecological weapon” that reshapes the local microbial community to facilitate pathogen colonization and persistence. Although 16S sequencing reveals compositional changes, additional studies are required to determine whether these effects result from direct inhibition of particular taxa, host-driven environmental selection, or a combination of both.

Our findings substantially extend the ecological framework proposed for holomycin in *Photobacterium galatheae*. In that system, holomycin was shown to promote biofilm formation without acting as a direct toxic factor^24^. Our study extends this phenomenon into an infection relevant cold-temperature (16℃) and reveals that the absence of holomycin decreased the biofilm formation, while its supplementation in Yru_*Δhom7* significantly restored biofilm formation in a dose-dependent manner (Figure 6). Mechanistically, holomycin may promote biofilm formation through its reactive disulfide bond facilitating extracellular matrix protein assembly^24,50-52^, which potentially enables quorum sensing activation and pathogenicity virulence factor delivery. Consistent with this, splenic transcriptomic analysis revealed that Yru_WT infection was associated with altered expression of host genes involved in innate immune recognition, adaptive immune priming, ubiquitin-like modification, and apoptosis, whereas these responses were markedly attenuated in the absence of holomycin (Figure 5). These processes are the targets of the main virulence factor-T3SS effector proteins, including YopJ-mediated NF-κB inhibition and YopH-mediated phagocytosis blockade^53-55^, suggesting a link between holomycin-dependent colonization and effector-mediated immune modulation. In addition, our data raise the possibility that holomycin may influence infection through mechanisms beyond biofilm formation. Our *Y. ruckeri* transcriptome revealed that holomycin biosnythesis is significanlty associated with the expression of virulence factor encoding genes (Figure 3E). Moreover, in Yru_*Δhom7*-infected fish the minimal splenic immune perturbation was observed at 24 h, when biofilm is unlikely to be fully established, suggests holomycin may contribute to infection through directly influencing the expression or delivery of virulence factors. Further investigation of virulence-associated pathways, such as T3SS or other colonization factors such as flagella and pili^56^, between Yru_WT and Yru_*Δhom7* will be necessary to clarify these mechanisms. Nevertheless, our data revealed holomycin as a central mediator linking biofilm formation, immune modulation, and pathogenicity in *Y. ruckeri* during fish infection.

To conclude, our work expands the ecological role of natural products from a producer-centred trait to a host-associated, microbiome-mediated strategy that shapes the outcome of host-pathogen interactions. By revealing how holomycin promotes infection through indirect ecological mechanisms rather than direct toxicity, this study provides a conceptual framework for targeting microbial competition and community dynamics as a means of disease control. These insights may inform the development of novel intervention strategies, including antivirulence therapies, microbiome-based management, and effective vaccine design, particularly for cold-water fish species that are highly vulnerable to *Y. ruckeri* infection. More broadly, our findings highlight the potential of integrating chemical ecology with host-microbiota interactions to improve disease prevention and sustainable aquaculture practices.

## RESOURCE AVAILABILITY

The authors declare that the data and materials supporting the findings reported in this study are available from the authors upon reasonable request.

## ACKNOWLEDGMENTS

This work was supported by the National Natural Science Foundation of China (32170079 to Z.Q., 32200035 to H.Z., 32400235 to J.H., and 22506007 to T.Z.), Guangdong S&T Program (2024B1111130001 to Z.Q.), the Natural Science Foundation of Guangdong (2024A1515012593 to Z.Q., 2023A1515110175 to J.H., and 2026A1515011616 to H.Z.), Guangdong Talent Scheme (2021QN020100 to Z.Q.). The authors would like to thank the Interdisciplinary Intelligence Super Computer Center, Beijing Normal University, for high performance computing for access to computational resources. We are also grateful to Prof. Peng Luo for kindly providing the *Yersinia ruckeri* genetic manipulation vector systems.

