## supplementary information for "Holomycin from *Yersinia ruckeri* mediates fish infection at low temperature"

**Table of contents**

**1. Experimental:**

**1a.** Standard chemistry methods and materials.

**1b.** Genome sequencing, assembly, and functional annotation of Yru_WT

**1c.** Phylogenetic analysis and evolutionary comparison of holomycin BGCs.

**1d.** Transcriptomic and proteomic integrative analysis of Yru_WT under different media.

**1e.** Scanning electron microscopy of Yru_WT and Yru_*Δhom7*.

**1f.** Production and purification of holomycin.

**1g.** In vivo pathogenicity and holomycin intervention in rainbow trout.

**1h.** Sample preparation and extraction of holomycin from bacterial cultures and rainbow trout tissues.

**1i.** RNA extraction and RT-qPCR.

**1j.** Histology examination of rainbow trout liver and spleens.

**1k.** Intestinal Microbiome Analysis via 16S rRNA Sequencing.

**1l.** RNA sequencing (RNA-seq) analysis of rainbow trout spleen.

**1m.** The formation of *Y. ruckeri* biofilms.

**1n.** Statistical analysis.

**2. ESI Figures**

**Figure S1.** Scanning electron microscopy images of Yru_WT and Yru_*Δhom7*.

**Figure S2.** Standard curve of holomycin.

**Figure S3.** $^1$H NMR spectrum of holomycin.

**Figure S4.** Representative H&E-stained liver, intestine, and gill sections from from fish in each group at 72 h post-infection.

**Figure S5.** Results of 16S rRNA sequencing.

**3. ESI Tables**

**Table S1.** Pairwise PERMANOVA results of β-diversity based on unweighted UniFrac distance. $R^2$ values indicate the proportion of variance explained by each comparison. $P$<0.05 were considered statistically significant.

**Table S2.** Strains, plasmids and primers used in this work.

**4. ESI References**

### 1. Experimental:

**1a. Standard chemistry methods and materials**. Acetonitrile (catalog A742210, ≥ 99.9% purity) and methanol (catalog M813904, ≥ 99.9% purity) of chromatographic grade were obtained from Macklin Biochemical Co., Ltd. (Shanghai, China). Ethyl acetate (catalog 141786, 99.5% purity) was obtained from Lingfeng Chemical Reagent Co., Ltd. (Shanghai, China). The Adenosine 5'-triphosphate disodium salt hydrate, PBS, Tris-(2-carboxyethyl)-phosphine hydrochloride, 2,6-Diaminoheptanedioic acid, Imidazole, Tris, isopropyl β-D-thiogalactopyranoside (IPTG), L-Cysteine, Magnesium chloride, Lysozyme, and L-(+)-Arabinose were obtained from Sangon Biotech Co., Ltd. (Shanghai, China).

**1b. Genome sequencing, assembly, and functional annotation of Yru_WT.** Yru_WT strain was cultured in tryptic soy broth (TSB) at 16°C with 180 rpm for 2 days. 1 mL of the bacterial solution was collected and centrifuged at 5000 rpm for 10 min. Cell pellets were washed three times with PBS. The total DNA was extracted and purified according to our previous study. DNA concentration and purity were measured using a NanoDrop 2000 spectrophotometer (Thermo Fisher Scientific, USA). Qualified DNA samples were sequenced using short-read (Illumina NovaSeq) and long-read (Oxford Nanopore MinION with R9.4.1 flow cell) platforms. Quality control, genome assembly, and the annotation of biosynthetic gene clusters (BGCs) for natural products were performed as described previously, with BGC analysis specifically identified using antiSMASH v7.0[1].

**1c. Phylogenetic analysis and evolutionary comparison of holomycin BGCs.** Genome sequences used in this study were extracted from NCBI using the relevant accession numbers and uploaded to the clinker online platform for holomycin BGC synteny analysis and gene annotation. Clusters were aligned based on homologous genes. Gene annotations and synteny relationships were visualized to evaluate conservation, rearrangements, and evolutionary divergence across strains.

**1d. Transcriptomic and proteomic integrative analysis of Yru_WT under different media.** Yru_WT was cultured in TSB (produce holomycin) and MRS (not produce holomycin) medium at 16°C with 180 rpm for 2 days. Cells were washed three times with PBS buffer and harvested by centrifugation at 5000 rpm for 10 min. For transcriptomic analysis, the total RNA was extracted by a bacterial kit (catalog DP430, TIANGEN, China) and purity were assessed by a Nanodrop 2000 spectrophotometer

(Thermo Scientific, USA). RNA sequencing was conducted by OE biotech and clean reads were aligned to the Yru_WT genome using HISAT2 v2.2.1[2]. Differential expression analysis was performed on the quantified reads using the DESeq2 R package, and differentially expressed genes (DEGs) were identified based on a cutoff of |log2FC| > 1. For proteomic analysis, cells were frozen in liquid nitrogen and analyzed by LC-MS/MS-based proteomics (OE Biotech). The differentially expressed proteins (DEPs) were defined based on a cutoff of |log2FC| >1.

**1e. Scanning electron microscopy of Yru_WT and Yru_*Δhom7*.** Yru_WT and Yru_ *Δhom7* strains were harvested by centrifugation at 5,000 rpm for 5 min and washed with PBS. The collected samples were fixed with 2.5% glutaraldehyde in PBS at 4°C overnight. Following fixation, the cells were dehydrated through a graded ethanol series (30%, 50%, 70%, 80%, 95%, and 100%) for 15 min at each concentration. The dehydrated samples were mounted onto clean silicon slices, followed by natural drying in the air, and then coated by a gold sputter coater. The surface morphology of the bacterial cells was examined using a field-emission scanning electron microscope (Hitachi Regulus 8100).

**1f. Production, purification and quantification of holomycin.**

Yru_WT was cultured in 40 L of TSB medium at 16°C with 180 rpm agitation for 4 days. The fermentation broth was extracted three times with an equal volume of ethyl acetate. The organic phases were pooled and concentrated to yield 3.39 g of crude extract. This extract was resuspended in 4 mL of ethyl acetate, mixed thoroughly with 5 g of silica gel (200-300 mesh size), and evaporated to dryness for dry loading. The mixed sample was loaded onto a chromatography column packed with normal-phase silica gel (200-300 mesh size) and eluted with dichloromethane-methanol. A total of 44 fractions were collected, and microbes' metabolites were analyzed using the LC-MS/MS method as described in 1e. Fractions containing holomycin were isolated and further purified using an Agilent 1290 series HPLC system under UV absorbance at 386 nm using the Phenomenex Kinetex C18 column (150 × 10 mm, 110 Å). The mobile phase contained 0.1% formic acid-water (A) and 0.1% formic acid-acetonitrile (B), following the linear gradient method: 0-5 min, 30% B; 5-30 min, 30%-70% B; 30-32 min, 70%-100% B; 32-42 min, 100% B; 8.1-9 min, 5% B with a constant flow rate of 2.5 mL/min. The injection volume was 18 µL.

**1g. *In vivo* pathogenicity and holomycin intervention in Rainbow Trout.** The rainbow trout (15 ± 1 g) were purchased from a commercial company in Shan dong province, China, and kept in tanks with aerated water at 16 ± 1°C. Rainbow trout were used to evaluate the pathogenicity of *Yru*_WT and the *in vivo* role of holomycin. including fish growth, tissue morphology, and immune responses. Intraperitoneal (IP) injection was selected as the administration route for both *Y. ruckeri* and holomycin to ensure efficient systemic absorption, as commonly applied in fish infection models. Based on our preliminary dose-response data, a bacterial density of $10^7$ CFU/L was utilized for all the bacteria infection assays. Rainbow trout were randomly assigned into different treatment groups and were treated with 50 µL 1 x PBS (N=35), Yru_WT (N=35), or Yru_*Δhom7* (N=35) through a single IP injection. At 24, 72, and 120 h post-injection, fish were euthanized with an overdose of MS-222. Livers and spleens were collected for histology, RT-qPCR, and RNA-sequencing. Intestines were collected for 16S rRNA gene sequencing, and whole-fish samples were preserved for the detection of holomycin. The mass spectrometry detection method for the compound refers to 1l.

To further elucidate the modulator role of holomycin, rainbow trout were treated by a single IP injection with 1 x PBS (N=40), Yru_WT (N=40), Yru_*Δhom7* (N=40), holomycin alone (~3µg / fish with average body weight of 15g, N=40), or a combination of Yru_*Δhom7* and holomycin (N=40). The holomycin dose was selected based on its *in vivo* production levels, adjusted by a magnification factor of 10 to account for the metabolism and excretion rates. The mortality rate of fish was recorded every 6 h post-injection. The five groups above were respectively injected with five fish and livers and spleens were collected for RT-qPCR at 72 h post-injection. Tissue lesions were assessed by observing hyperemia area. All animals were maintained and treated according to the *Guide for the Care and Use of Laboratory Animals* and approved by the *Institutional Animal Care and Use Committee protocol* of Shanghai Ocean University (Approval No. SHOU-DW-2021-027).

**1h. Sample preparation and extraction of holomycin from rainbow trout.** The homogenized rainbow trout infected with 1 x PBS (N=5), Yru_WT (N=5), and Yru_*Δhom7* (N=5) were extracted with ethyl acetate at a ratio of 1:8. The mixtures were vortexed, sonicated for 15 min, and soaked in ethyl acetate for 12 h. The supernatant of mixture was collected and evaporated to dryness under a rotary evaporator. The extracts were reconstituted in 300 µL methanol and centrifuged at 15000 rpm for 20 min. Holomycin in supernatant was analyzed by UPLC-MS/MS method as described in 1e.

**1i. RNA extraction and RT-qPCR.** Spleens were collected from rainbow trout infected with 1 x PBS (N=6), Yru_WT (N=6), and Yru_*Δhom7* (N=6) to examine the mRNA expression of immune-related genes. Total RNA of spleen tissues was extracted using the TRIzol reagent (catalog 15596026, Invitrogen, CA, USA) according to the manufacturer's instructions. RNA concentration and purity were measured using NanoDrop. Total RNA was then reverse transcribed into cDNA using the HiScript III RT SuperMix (catalog R323, Vazyme) and stored at -80°C. The RT-qPCR was performed in 96-well plates using the SYBR Green system (catalog Q711, Vazyme) in an Applied Biosystems 7500 Fast real-time PCR system (CFX96, Bio-Rad, USA). RT-qPCR thermocycle was programmed for initial denaturation at 95°C for 30 s, 40 cycles of 95 °C for 10 s and 60 °C for 30 s, followed by a melt-curve analysis. The relative expression levels of immune-related genes, including *IL-1β*, *IL-6*, *IL-8*, and *IL-10* were calculated using the $2^{-\Delta\Delta Ct}$ method and normalized to the expression of the housekeeping gene EF-1α. The primer sequences of all examined genes are listed in Supplementary Table S1. Each experiment was performed with three biological replicates in technical triplicate.

**1j. Histology of rainbow trout livers and spleens.** Liver and spleen tissues were collected from rainbow trout infected with PBS (N=5), Yru_WT (N=5), and Yru_*Δhom7* (N=5), and immediately fixed in 10% neutral buffered formalin overnight. Following fixation, the tissues were dehydrated through a graded series of ethanol, cleared in xylene, and embedded in paraffin. The embedded tissues were then serially sectioned at a thickness of 4 micrometers using microtome. The sections were stained with hematoxylin and eosin (H&E, Thermo Fisher Scientific) and imaged using microscope (ZEISS Axio Imager Z2). Histopathological changes in livers and spleens from fish were examined, including tissue necrosis, inflammatory cell infiltration, and structural alterations.

**1k. Intestinal Microbiome Analysis via 16S rRNA Sequencing.** Intestine were collected from the rainbow trout treated with 1xPBS (N=5), Yru_WT (N=5), and Yru_*Δhom7* (N=5) at 72 h-post IP injection. Total genomic DNA was extracted using the MagPure Soil DNA LQ Kit (Magan). DNA concentration and intergrity was assessed by NanoDrop 2000 and agarose gel electrophoresis. The [V3-V4] region was amplified with barcoded primers [343F/798R] using Takara Ex Taq (catalog HRR001A, Takara, Japan). Libraries were purified with AMPure XP beads (Agencourt), quantified using a Qubit dsDNA Assay Kit (catalog Q32854, Thermo Fisher Scientific), and sequenced on an Illumina NovaSeq 6000 (2 × 250 bp) (Illumina Inc., USA; OE Biotech Company, China). Reads were trimmed using Cutadapt software and processed in QIIME2 (v2020.11) using DADA2 for quality filtering, denoising, merging, and chimera removal to obtain the representative read of each ASV. Taxonomic assignment was performed using SILVA v138 with q2-feature-classifier. The microbial diversity was estimated using the alpha diversity that include Chao1 and Shannon index. The unweighted Unifrac distance matrix was used for unweighted Unifrac Principal coordinates analysis (PCoA) to estimate the beta diversity. The linear discriminant analysis effect size (LEfSe) was used to identify differentially abundant taxa.

**1l. RNA sequencing (RNA-seq) analysis of Rainbow Trout spleens.** Spleens were collected from the Rainbow Trout treated with 1xPBS (N=3), Yru_WT (N=3), and Yru_*Δhom7* (N=3) at 72 h post IP injection. Total RNA was extracted using the TRIzol reagent (Invitrogen). Quality and quantity of RNA were evaluated with Nanodrop 2000 spectrophotometer (Thermo Fisher Scientific). cDNA synthesis and library preparation were performed using the VAHTS Universal V10 RNA-seq Library Prep Kit (Premixed Version). Sequencing was conducted on an an Illumina Novaseq X Plus platform and 150 bp paired-end reads were generated. Raw reads of fastq format were firstly processed using fastp and the low-quality reads were removed to obtain the clean reads. High-quality trimmed paired sequencing reads were mapped to the reference genome using HISAT2. FPKM of each gene was calculated and the read counts of each gene were obtained by HTSeq-count. Principle component analysis (PCA) was performed using R PCAtools to evaluate the biological duplication of samples. Genes with an absolute FC > 2.0 and a false discovery rate (FDR) adjusted p-value < 0.05 were defined as significantly differentially expressed genes (DEGs). Gene ontology (GO) analyses and Kyoto Encyclopedia of Genes and Genomes (KEGG) pathway enrichment analysis were performed using Database for Annotation, Visualization, and Integrated Discovery (DAVID).

**1m. The formation of *Y. ruckeri* biofilms**. The biofilm formation ability of Yru_WT and Yru_*Δhom7* was assessed using the crystal violet staining method. Briefly, Yru_WT and Yru_*Δhom7* were cultured at 16°C for 12 h and diluted to an $OD_{600}$ of 0.3 in TSB. Then, bacteria were cultured in 24-well plates at 16°C for 3 days. In the complementation experiments, holomycin was added to cultures from 2 µM to 30 µM after 10 h of incubation. The bacteria were removed and wells were washed with Milli-Q water for three times. Biofilms in each well were stained with 1% crystal violet for 15 min. The crystal violet was removed and washed three times with water. The wells were dried for 30 min, added with 95% ethanol, and incubated for 20 min to dissolve the staining color. The solubilized crystal violet mixture was divided into three equal parts in a 96-well plate, and absorbance was measured at 590 nm.

**1n. Statistical analysis.** Statistical analysis was performed using GraphPad Prism version 9 (GraphPad Software, San Diego, CA). Student's t-test was performed to compare the numerical data of two treatment groups. One-way ANOVA followed by a Tukey's multiple comparison test and 'Dunnett' adjustment was used to compare numerical data of multiple treatment groups. The data was reported as the mean value ± standard deviation (SD). Statistical significance was defined as a p-value $< 0.05$.

### 2. ESI Figures

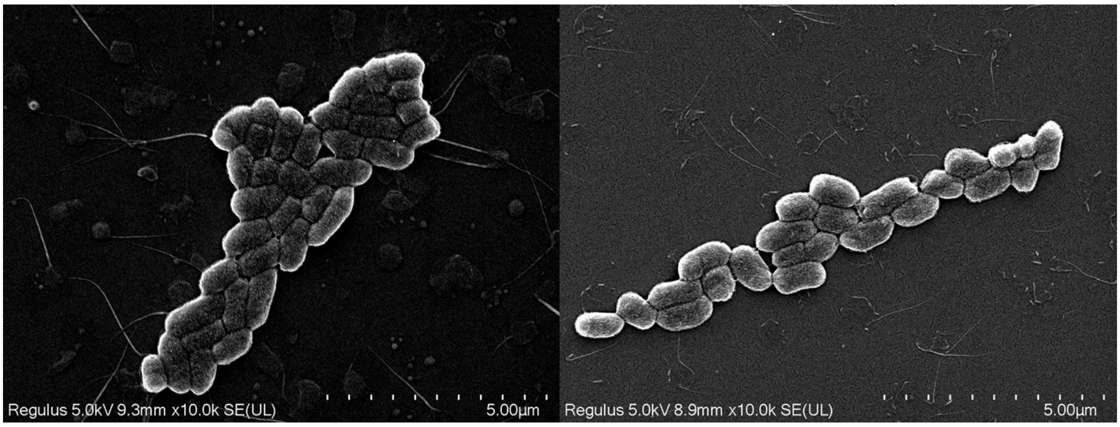


**Figure S1.** Scanning electron microscopy images of Yru_WT (left panel) and Yru_Δ*hom7* (right panel).

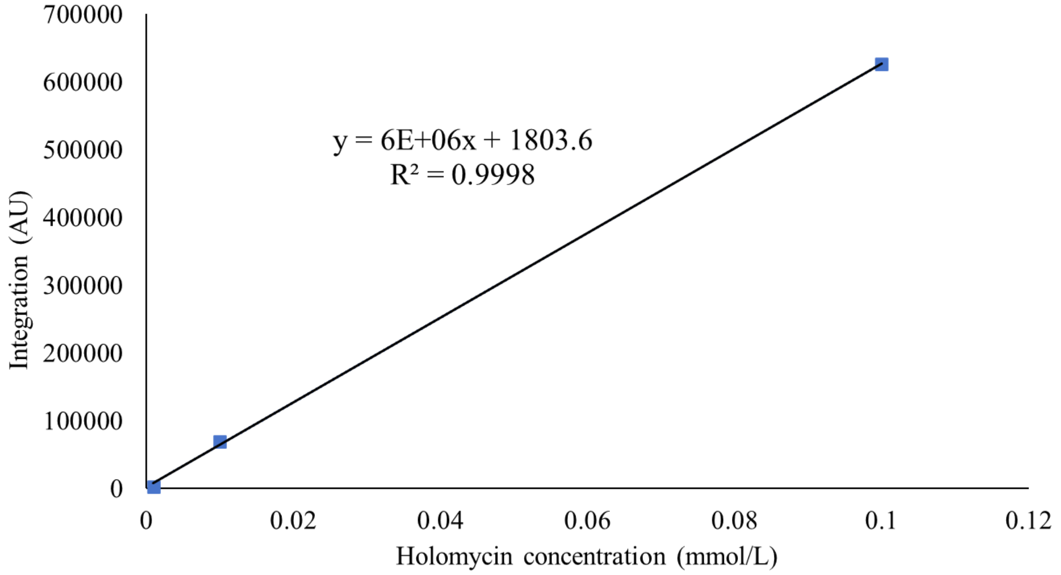


**Figure S2.** Standard calibration curve of holomycin. Each data point was shown as mean ± SD (n = 3).

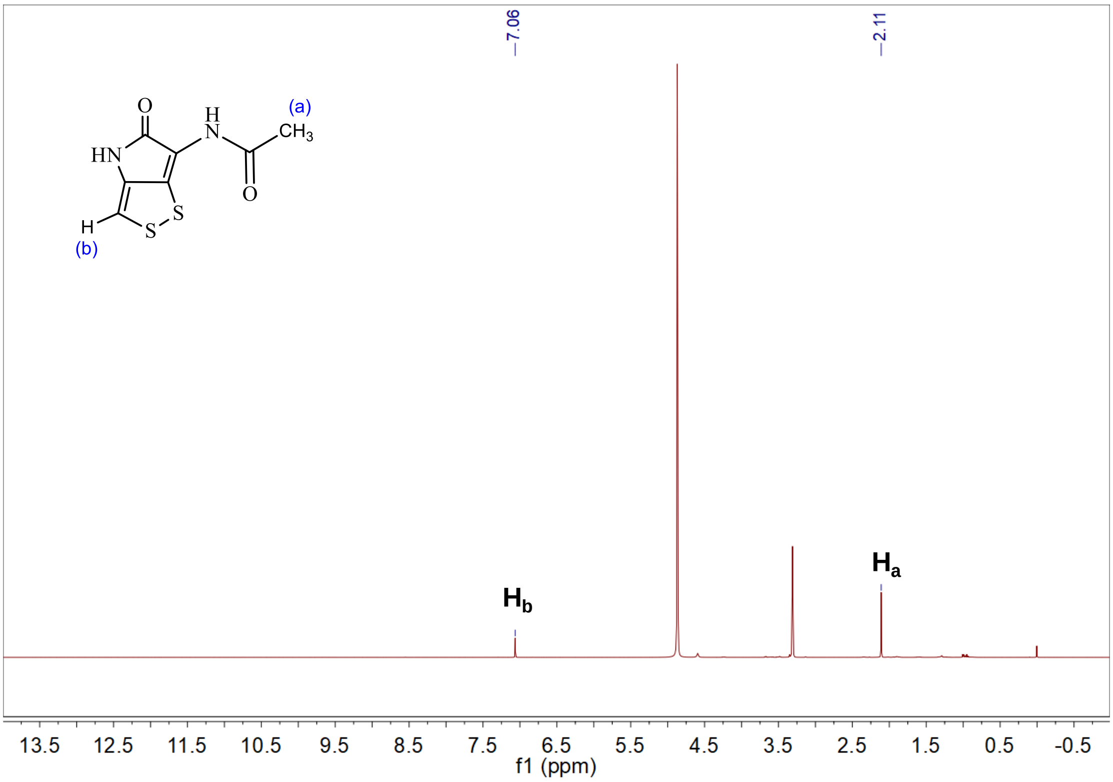


**Figure S3.** $^1$H NMR spectrum of holomycin, 400 MHz, methanol, $CD_3OD$.

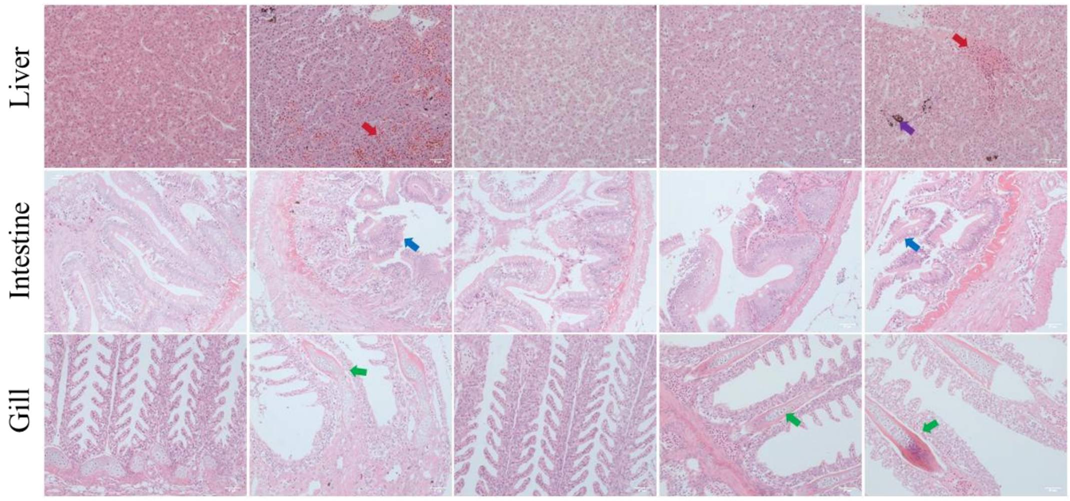


**Figure S4.** Representative H&E-stained liver, intestine, and gill sections from fish in each group at 72 h post-infection. Red arrows indicate haemorrhagic lesions; purple arrows indicate melanomacrophage centres; blue arrows indicate the disruption of the intestinal folds; green arrows indicate vacuole-like structures.

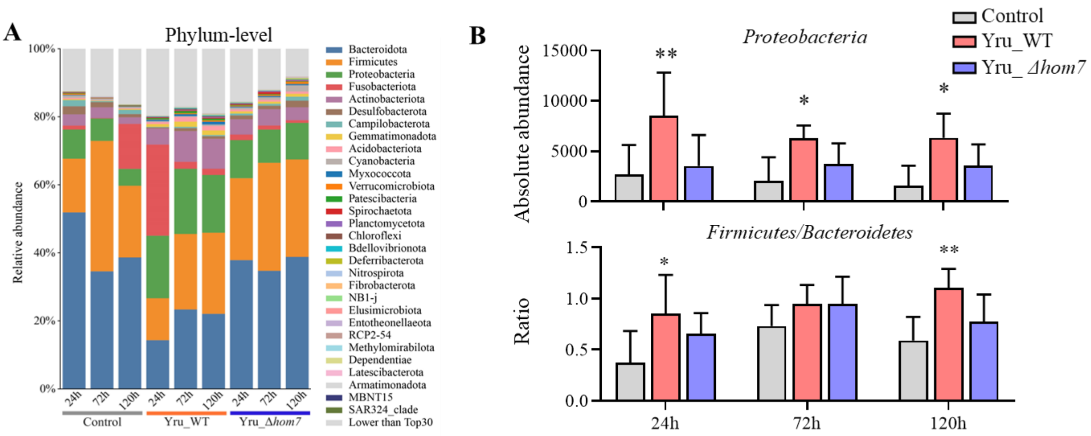


**Figure S5.** Results of 16S rRNA sequencing. (A) Phylum-level composition of rainbow trout gut microbiota at 72 h post-infection. Top 30 phyla are displayed. The remaining taxa are grouped as “Lower than Top30”. (B) Absolute abundance of *Bacteroidota* and the *Firmicutes*/*Bacteroidetes* ratio in fish infected with PBS, Yru_WT, or Yru_*Δhom7* at 24 h, 72 h, and 120 h post-infection.

### 3. ESI Tables

**Table S1.** Pairwise PERMANOVA results of β-diversity based on unweighted UniFrac distance. $R^2$ values indicate the proportion of variance explained by each comparison. $P<0.05$ were considered statistically significant.

| Time | group | *R2* | PERMANOVA Pvalue |
|---|---|---|---|
| 24h | All groups | 0.20564 | 0.036 |
| 24h | Yru_WT *vs* PBS | 0.1918 | 0.026 |
| 24h | Yru_*Δhom7* *vs* PBS | 0.13874 | 0.186 |
| 24h | Yru_WT vs Yru_*Δhom7* | 0.15462 | 0.056 |
| 72h | All groups | 0.23755 | 0.004 |
| 72h | Yru_WT *vs* PBS | 0.24315 | 0.013 |
| 72h | Yru_*Δhom7* *vs* PBS | 0.18594 | 0.027 |
| 72h | Yru_WT *vs* Yru_*Δhom7* | 0.12965 | 0.019 |
| 120h | All groups | 0.22192 | 0.005 |
| 120h | Yru_WT *vs* PBS | 0.21052 | 0.021 |
| 120h | Yru_*Δhom7* *vs* PBS | 0.16112 | 0.066 |
| 120h | Yru_WT *vs* Yru_*Δhom7* | 0.15242 | 0.027 |

**Table S2.** Strains, plasmids and primers used in this work.Error! Not a valid link.

| **Primer** | **Sequence** | **usage** |
|---|---|---|
| *hom1*-qF | CACTTCTGCCCGGTTACCTC | qRT-PCR |
| *hom1*-qR | ATCAGCCCACAACCTCATGG | |
| *hom2*-qF | AGAACATGTCGCTATCTGG | |
| *hom2*-qR | GAGATTCACCTCGTCAGCA | |
| *hom3*-qF | ACGGAAGGCGCATCAGTAAA | |
| *hom3*-qR | TCAAGCGTCACCAGAGCATT | |
| *hom4*-qF | TGCTGGTCATCGAACGTGAT | |
| *hom4*-qR | CCTTGGCATACCGCCCATAA | |
| *hom5*-qF | GGCATCACAGAGTTTGCCAC | |
| *hom5*-qR | CGTCAGACCACAAGTAGCGA | |
| *hom6*-qF | GCTTATCTCACTCCAGCC | |
| *hom6*-qR | ACACCAGAACGCTGGAGAAG | |
| *hom7*-qF | TCAGGATTTCACGCTCACC | |
| *hom7*-qR | CGAACGAGTCAGAGGATGGG | |
| *hom15*-qF | AGGTTTCGGTTTCATCACTCCA | |
| *hom15*-qR | CGAACTCTACGTTCTGGCCTT | |
| 16S-qF | AAGAAGCACCGGCTAACTCC | |
| 16S-qR | GTTAAGCGCGGGGATTTCAC | |
| Plp12-F | GACATATCGTCGCTAGCTGTTCC | Plp12 plasmid backbone amplificatio n |
| Plp12-R | CGACTCAAGGTCTAGATTCCTGC | |
| *hom7*-UF | GAATCTAGACCTTGAGTCGGTCAGTCCAGAG CTTGGAAT | Amplificati on of HAL and HAR for deletion of *hom7* |
| *hom7*-UR | GGACAATCCCATATCCTGACAATTCACTCCAC TCATCTGTG | |
| *hom7*-DF | CACAGATGAGTGGAGTGAATTGTCAGGATAT GGGATTGTCC | |
| *hom7*-DR | CAGCTAGCGACGATATGTCAAGGAATGCCGT GACTGACG | |
| *hom7*-NF | ATCCTGACGTTCTATTGTGAG | Verification of the gene deletion of *hom7* |
| *hom7*-NR | ATGATCTGAAGATTACTGCGC | |
| pBAD33-F | CTAGAGTCGACCTGCAGGCA | pBAD33 plasmid backbone amplificatio n |
| pBAD33-R | AGCTCGAATTCGCTAGCCCA | |

| | | |
|---|---|---|
| *hom7*-pBAD33-F | TGGGCTAGCGAATTCGAGCTATGAAAATGATGCGGGAAGCTACC | Cloned *hom7* for complementation |
| *hom7*-pBAD33-R | TGCCTGCAGGTCGACTCTAGTCATGACAAGGCTTCCTCAGGATC | |

### 4. ESI References


1. Huang, J., Wu, Y., Gao, Q., Li, X., Zeng, Y., Guo, Y., Zhang, H., and Qin, Z. (2024). Metagenomic exploration of the rhizosphere soil microbial community and their significance in facilitating the development of wild-simulated ginseng. Applied and Environmental Microbiology *90*, e02335-02323. doi:10.1128/aem.02335-23.
2. Kim, D., Paggi, J.M., Park, C., Bennett, C., and Salzberg, S.L. (2019). Graph-based genome alignment and genotyping with HISAT2 and HISAT-genotype. Nature Biotechnology *37*, 907-915. 10.1038/s41587-019-0201-4.
